# Up, down, and out: optimized libraries for CRISPRa, CRISPRi, and CRISPR-knockout genetic screens

**DOI:** 10.1101/356626

**Authors:** Kendall R Sanson, Ruth E Hanna, Mudra Hegde, Katherine F Donovan, Christine Strand, Meagan E Sullender, Emma W Vaimberg, Amy Goodale, David E Root, Federica Piccioni, John G Doench

## Abstract

Advances in CRISPR-Cas9 technology have enabled the flexible modulation of gene expression at large scale. In particular, the creation of genome-wide libraries for CRISPR knockout (CRISPRko), CRISPR interference (CRISPRi), and CRISPR activation (CRISPRa) has allowed gene function to be systematically interrogated. Here, we evaluate numerous CRISPRko libraries and show that our recently-described CRISPRko library (Brunello) is more effective than previously published libraries at distinguishing essential and non-essential genes, providing approximately the same perturbation-level performance improvement over GeCKO libraries as GeCKO provided over RNAi. Additionally, we developed genome-wide libraries for CRISPRi (Dolcetto) and CRISPRa (Calabrese). Negative selection screens showed that Dolcetto substantially outperforms existing CRISPRi libraries with fewer sgRNAs per gene and achieves comparable performance to CRISPRko in the detection of gold-standard essential genes. We also conducted positive selection CRISPRa screens and show that Calabrese outperforms the SAM library approach at detecting vemurafenib resistance genes. We further compare CRISPRa to genome-scale libraries of open reading frames (ORFs). Together, these libraries represent a suite of genome-wide tools to efficiently interrogate gene function with multiple modalities.tracr

## INTRODUCTION

Pooled genetic screening requires the reliable perturbation of a large number of targets across the genome. The robust on-target activity and high fidelity of the CRISPR-Cas9 system has allowed it to quickly surpass RNAi technology as the method of choice for genetic screening^1,2^. Moreover, unlike RNAi, the CRISPR technology can also be used to screen a variety of perturbations beyond gene down-regulation: unmodified Cas9 can generate complete loss-of-function alleles, and a nuclease-deactivated Cas9 (dCas9) may be tethered to an inhibitory or activating domain to regulate gene expression via CRISPR interference (CRISPRi) or CRISPR activation (CRISPRa), respectively^3-8^. In comparison to CRISPR knockout (CRISPRko), CRISPRi and CRISPRa allow for the transient modulation of gene expression, which can reveal novel phenotypes and enable more flexible experimental designs.

Although even early CRISPR libraries outperformed RNAi libraries, the selection of highly active sgRNAs can further improve library performance^9-11^. Numerous approaches have been developed to predict sgRNA activity based on sequence features, position, and off-target activity, which have been recently summarized^12^. These prediction algorithms have been used to develop next-generation genome-wide CRISPRko libraries for pooled screening^13-17^. Similarly, selection criteria have been used to develop genome-wide libraries for CRISPRi^18,19^ and CRISPRa^18-20^. Some libraries contain many sgRNAs per gene, allowing hits to be detected with high confidence. However, screening efficiency can also be improved by looking at libraries more holistically; well-designed libraries that effectively modulate targeted genes with fewer sgRNAs per gene provide more information with less investment of resources. This is particularly useful in settings where cell numbers are limiting, such as screens in primary cells or *in vivo.* Here, we evaluate existing genome-wide libraries for CRISPRko, CRISPRi, and CRISPRa with S. *pyogenes* Cas9 (SpCas9) via 16 screens conducted across 3 cell lines. We present the dAUC (delta area under the curve) metric, an intuitive and size-unbiased metric for library performance in negative selection, loss-of-function screens that quantifies the ability of a genome-wide library to distinguish essential and non-essential genes. We then provide data from negative selection screens with Brunello, our optimized CRISPRko library, and demonstrate that Brunello outperforms previously published CRISPRko libraries, on both the sgRNA and the gene level.

We also introduce two new genome-wide libraries: Dolcetto, a human CRISPRi library, and Calabrese, a human CRISPRa library. We demonstrate that Dolcetto outperforms existing CRISPRi libraries in negative selection screens and performs comparably to CRISPRko in depletion of essential genes. In positive selection screens, we show that Calabrese finds more vemurafenib resistance genes than the SAM library approach. We also compare CRISPRa to screens performed with open reading frame (ORF) overexpression libraries and show that these technologies identify a number of common genes, but also each uncover distinct and complementary hits. Our findings demonstrate the importance of optimized library design to improve the quality of genetic screens using CRISPRko, CRISPRi, and CRISPRa.

## RESULTS

### Brunello, a genome-wide CRISPRko library

We previously reported the design of optimized CRISPRko sgRNA libraries, Brunello and Brie, designed for improved on-target and reduced off-target activity in the human and mouse genomes, respectively^16^. The Brunello library comprises 77,441 sgRNAs, which includes an average of 4 sgRNAs per gene and 1,000 non-targeting control sgRNAs. We conducted genome-wide negative selection (dropout) screens in A375 (melanoma) and HT29 (colon cancer) cells that were first engineered to expressed Cas9. The Brunello library was cloned into the lentiGuide vector^2^ and transduced into cells in biological replicates at a multiplicity of infection (MOI) of ~0.5 and passaged at a minimum of 500x coverage; that is, the majority of transduced cells received only a single viral integrant and each sgRNA is present, on average, in 500 unique cells. Uninfected cells were removed with puromycin selection, and the population was cultured for a total of three weeks, after which genomic DNA was harvested, the sgRNA cassette retrieved by PCR, and the abundance of sgRNAs quantitated by Illumina sequencing. All pooled screens described herein follow these general experimental parameters.

To compare the performance of Brunello to other libraries, we developed a sgRNA-level metric assessing the performance of viability screens. The use of gold standard gene sets of 1,580 essential and 927 non-essential genes^15,21^ allows the calculation of the area under the curve (AUC) for all sgRNAs targeting essential and non-essential genes (Fig. 1a). In viability screens, the sgRNAs targeting essential genes should be among the most depleted compared to their starting abundance. A random distribution results in an AUC of 0.5; thus, an effective library should have an AUC > 0.5 for sgRNAs targeting essential genes, indicating that these sgRNAs preferentially deplete, and an AUC ≤ 0.5 for sgRNAs targeting non-essential genes and non-targeting sgRNAs, indicating that these sgRNAs preferentially remain. Importantly, the AUC metric is an sgRNA-level analysis that effectively assesses the quality of sgRNA design in a library. Because every sgRNA in the library is included in the AUC – rather than collapsing sgRNAs to genes or subsampling to include only the top-performing sgRNAs in a library – this metric is not biased towards larger libraries, which may achieve strong performance through sheer force of a large number of sgRNAs. The AUC is therefore a valuable metric to compare sgRNA design across libraries of different sizes.

**Figure 1.**
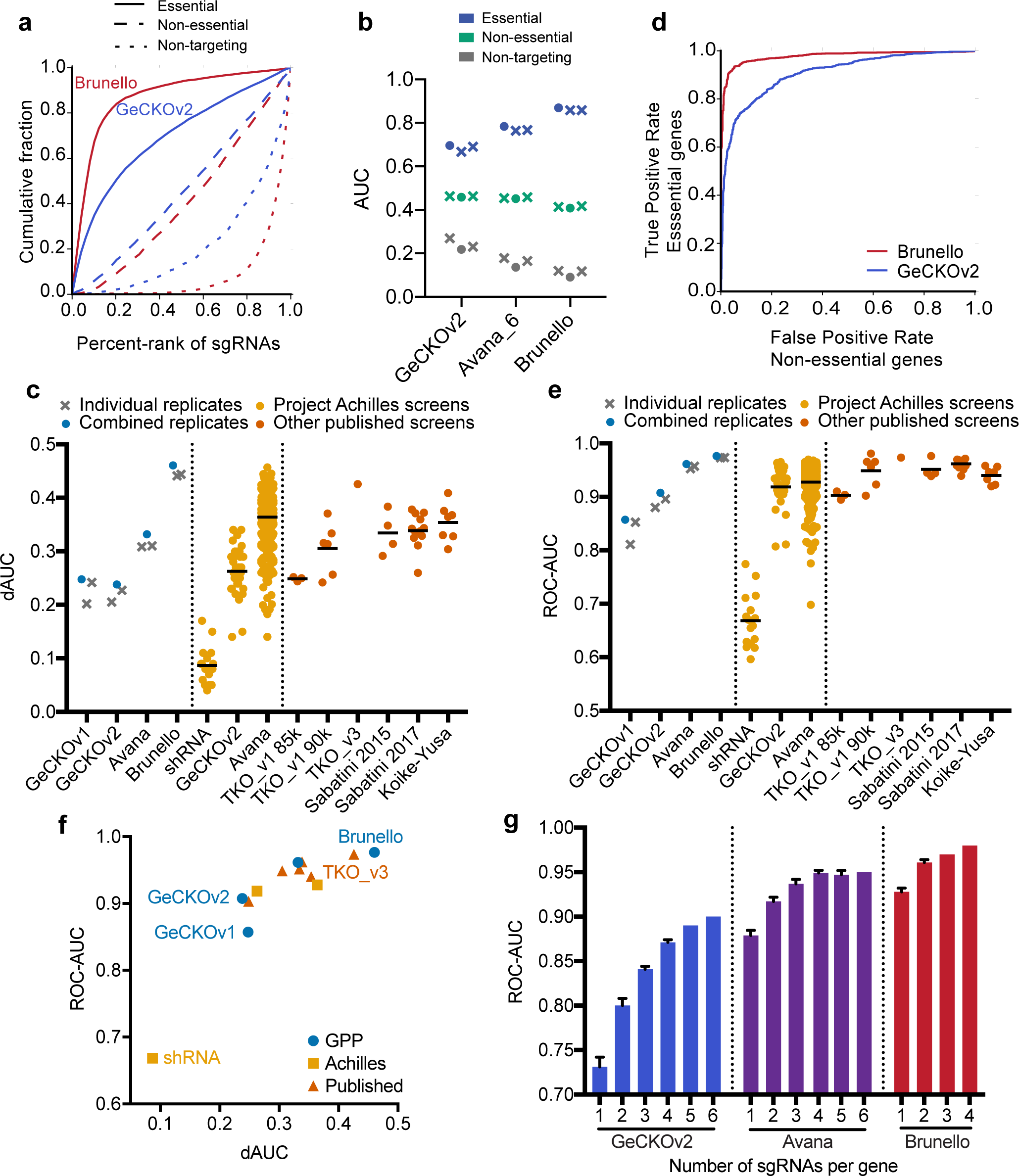
Improved performance of genome-wide CRISPRko libraries. (a) Area under the curve analysis of cell viability screening data for core essential (solid line), non-essential (dashed line) and non-targeting (dotted line) gene sets in the Brunello and GeCKOv2 library screened in A375 cells. (b) Comparison of AUCs for essential, non-essential and non-targeting sgRNAs across the generations of libraries screened by this group. The AUC of individual replicates are plotted as X’s, while the AUC values calculated from the averaged log2-fold change of the replicates are plotted as circular points. The Avana library was screened with all 6 sgRNAs per gene. (c) Comparison of the dAUC across different CRISPRko libraries. Individual replicates of the screens conducted by this group are plotted as X’s and the dAUC values calculated using the average log2-fold change of replicates are plotted as circular points. All other published library statistics are plotted as circular points from the combination of replicates, if provided. Black line represents the average of the dAUCs. For Project Achilles, a version of the Avana library with 4 sgRNAs per gene was used. (d) Receiver operating characteristic analysis of cell viability screening data for Brunello and GeCKOv2 library screened in A375 cells. False positive rates are calculated for non-essential genes and plotted against the true positive rate for essential genes. (e) Comparison of the ROC-AUC across different CRISPRko libraries. Values are plotted as in (c). (f) Scatter plot comparing dAUC and ROC-AUC values for different libraries; when multiple cell lines of data were available, the mean is plotted. GPP refers to screens executed herein and previously by this group. (g) Subsampling analysis calculating the ROC-AUC of n sgRNAs per gene in three different libraries. Error bars represent different iterations of library sampling.

In A375 cells, which were screened previously with the GeCKO and Avana libraries^2,16^, the Brunello library showed even greater depletion of sgRNAs targeting essential genes (AUC = 0.80; Fig 1b). As expected, sgRNAs targeting non-essential genes showed no evidence of depletion (AUC = 0.42). Conversely, non-targeting sgRNAs were among the least depleted, with an AUC = 0.16. That these sgRNAs are preferentially retained is expected based on the well-described cutting effect that manifests in CRISPRko screens, whereby dsDNA breaks caused by sgRNAs lead to a detectable effect on cell growth; in extreme cases, such as copy number amplified target sites or promiscuous sgRNAs, this effect is greatly magnified^16,22-24^.

To simplify comparisons across libraries, we next calculated the difference between the AUC of sgRNAs targeting essential and non-essential genes (delta AUC, dAUC). We analyzed previous screens using this metric and found that the dAUC increased with each generation of CRISPRko library design, from GeCKO to Avana to Brunello, validating that improvements in sgRNA and library design led to increased performance; merging of multiple replicates only led to a small increase in the dAUC, suggesting that individual replicates were well-powered (Fig 1c). Notably, the improvement in dAUC from GeCKOv2 to Brunello (delta dAUC, ddAUC = 0.46 - 0.24 = 0.22) was greater than the average improvement from RNAi to GeCKOv2 in Project Achilles^25^ (ddAUC = 0.26 - 0.09 = 0.17). We then calculated dAUCs for a number of screens using previously published CRISPRko libraries^10,14,15,17,23,24,26^ and found that Brunello outperformed other existing genome-wide libraries by this metric (Fig 1c). Another recent library, TKOv3, which was screened in the haploid cell line HAP1, was the next-best performer by this metric^10^. Importantly, the design rules underlying Brunello were not trained on data from negative selection screens^16^ and thus the dAUC metric represents an unbiased measurement of the performance of this library.

Next, to compare libraries at the gene level, we averaged all sgRNAs targeting a gene and performed precision-recall analysis, defining essential genes as true positives and non-essential genes as false positives, and calculated the area-under-the-curve of the receiver-operator characteristic (ROC-AUC) (Fig. 1d, e). Compared to the dAUC metric, the ROC-AUC metric shows the value of having more sgRNAs per gene; for example, GeCKOv1 and GeCKOv2 used the same design criteria and thus have similar dAUC values at the sgRNA level, but GeCKOv2, with 6 sgRNAs per gene, substantially outperformed GeCKOv1, with 3-4 sgRNAs per gene, via the ROC-AUC metric (Fig. 1f). Notably, Brunello contains only 4 sgRNAs per gene but outperformed all other libraries by both metrics, highlighting the substantial effect of sgRNA design on library performance. We note that a recent study performed a similar comparison of some of these libraries^11^, although that study considered only 8,948 total genes, and used a set of essential genes that were partially defined by the CRISPR libraries under evaluation^10^, which may result in a biased estimate of performance.

We next performed subsampling analysis of sgRNAs in the Brunello, Avana, and GeCKOv2 libraries. After random draws of *n* sgRNAs per gene, the log2-fold change of the sgRNAs targeting the same gene were averaged, and the ROC-AUC was calculated (Fig 1g). Even with only a single sgRNA per gene, the Brunello library outperformed the GeCKOv2 library with the full 6 sgRNAs per gene, again demonstrating the large improvement in sgRNA design.

Additionally, we observed only a minor increase in performance with additional sgRNAs for the Brunello library, suggesting that libraries consisting of only 2 or 3 sgRNAs per gene can still be expected to perform well, an especially important criterion when cell numbers are limiting.

### Modifications to the tracrRNA

While sgRNA design is critical to maximizing on-target activity, other components of the CRISPR technology have also been optimized. Previously, several modifications to the tracrRNA have been proposed to increase on-target activity, including the removal of a potential U6 Pol III termination site and the extension of the first tracrRNA-derived stem loop to enhance the assembly of the sgRNA-Cas9 complex^27^. More detailed analyses further showed that the use of a T-to-C and compensatory A-to-G substitution provided the best activity, and that a five nucleotide extension of the stem was optimal^28^. A tracrRNA with these modifications was suggested to improve the on-target activity of sgRNAs in a CRISPRko screen with a focused test library^29^. However, we sought to assess the performance of a modified tracrRNA in a genome-wide setting.

We designed a modified tracrRNA for use in lentiCRISPRv2, hereafter referred to as tracr-v2, which removed the Pol III termination site and extended the first stem loop by 5 base pairs (Fig. 2a). To test on- and off-target activity with tracr-v2, we designed a tiling library containing all possible sgRNAs targeting EEF2, a core essential gene, with both perfect match and single-mismatched sgRNAs; notably, this latter class was not assessed previously^29^. We performed a negative selection screen in A375 cells, tiling across the essential gene EEF2, with sgRNAs targeting CD81 serving as controls, as we had done previously with a focused library to compare S. *aureus* Cas9 (SaCas9) to SpCas9^30^. Consistent with other reports, perfect match sgRNAs with tracr-v2 showed significantly stronger depletion of EEF2 than the original tracrRNA, indicating improved on-target activity (Fig. 2b). However, tracr-v2 also exhibited higher levels of off-target activity with mismatched sgRNAs (Fig. 2b).

**Figure 2.**
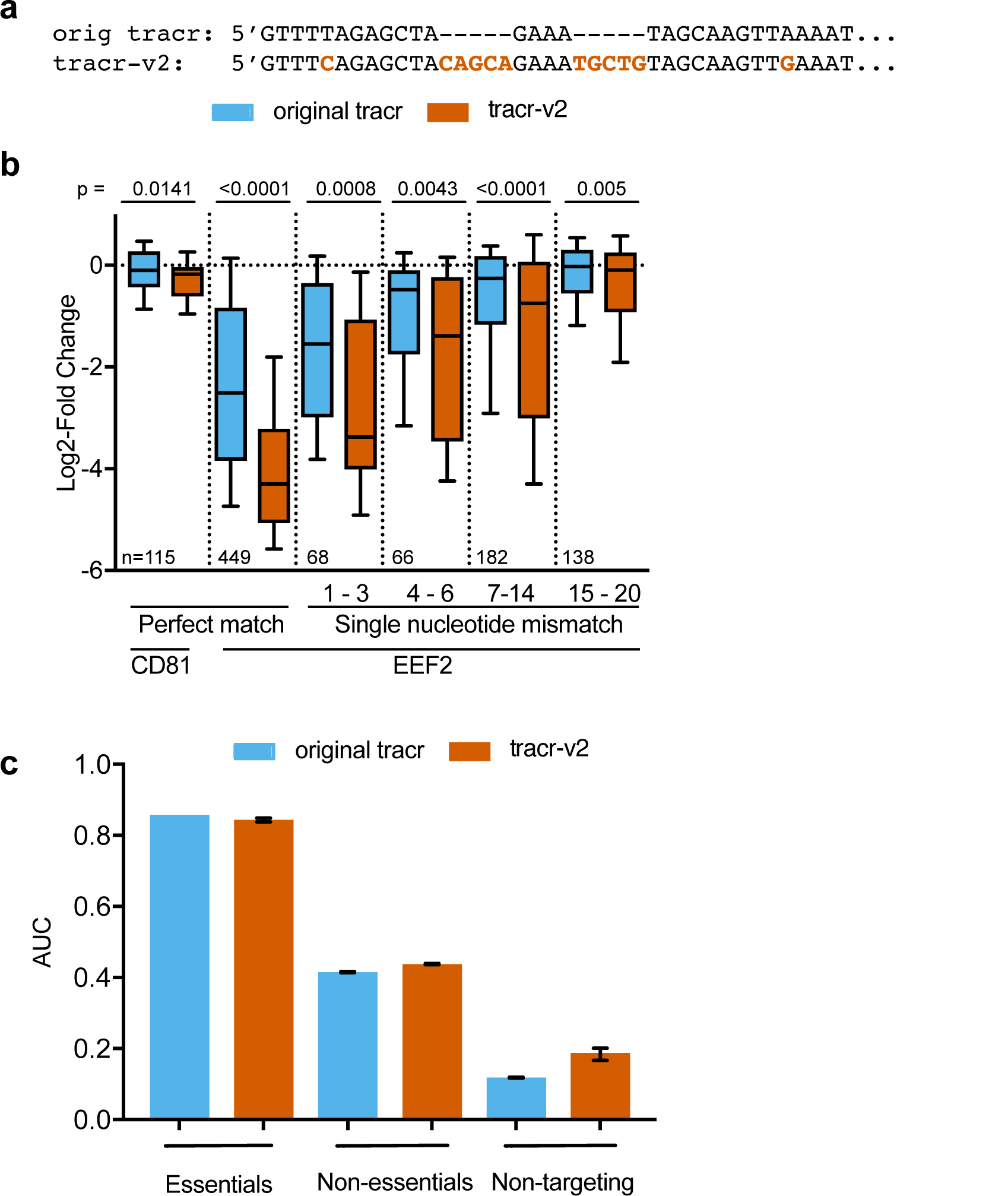
Evaluation of an alternative tracrRNA. (a) Comparison of the sequence of the original tracrRNA and tracr-v2. (b) Comparison of the log2-fold-change of perfectly matched sgRNAs and single base mismatches targeting the essential gene EEF2 for the original tracrRNA and tracr-v2. CD81-targeting sgRNAs serve as the control. Mismatched sgRNAs are binned by position, and are numbered such that nt 20 is PAM-proximal. The box represents the 25th, 50th and 75th percentiles, and whiskers show 10th and 90th percentiles. Two-tailed Welch’s t-test was used to compare the distributions within each bin and p values are indicated. The number of sgRNAs in each comparison is shown at the bottom. (c) AUC values for essential, non-essential, and non-targeting gene sets with the original tracrRNA and tracr-v2 in genome-wide dropout screens. Error bars represent the range of two biological replicates.

To test how this increase in both on- and off-target activity would affect performance of a genome-wide screen, we performed a negative selection screen using Brunello with tracr-v2 in lentiCRISPRv2. We observed that Brunello with tracr-v2 performed similarly to our previous results with the original tracrRNA (Fig 2c); tracr-v2 had a dAUC of 0.42, compared to 0.46 with the original tracrRNA. These results suggest that tracr-v2 neither improves nor harms library performance in CRISPRko genome-wide screens, at least in the case of a library selected to have active sgRNAs; we cannot rule out that a modified tracrRNA structure would have more beneficial impact on a library with less on-target activity, such as GeCKOv2.

### Dolcetto, a genome-wide CRISPRi library

Although CRISPRko screens have been widely employed and can identify hit genes with strong statistical significance, the technology has several limitations that require consideration. First, because the gene dosage is complete knockout in many cases, positive selection screens may miss genes that are necessary for viability at some level but show a phenotype upon partial inhibition. Second, genes with an increased copy number show a strong cutting effect when targeted with CRISPRko: although they may not be essential, the large number of dsDNA breaks has a strong viability effect that can mimic gene essentiality^22-24^.

A useful orthogonal approach to study loss-of-function phenotypes is CRISPR interference (CRISPRi). Here, point mutations engineered into Cas9 inactivate the nuclease domains, creating an RNA-guided DNA binding protein (dCas9), which can then be tethered to repressive domains such as KRAB to prevent efficient transcription^4,7,8,18^. Although a large portion of a protein coding gene can serve as an effective target for CRISPRko, CRISPRi is effective over a much more narrow region, near the transcription start site (TSS) of a gene. Previous studies have shown that the FANTOM database^31^, which relies on Capped Analysis of Gene Expression (CAGE) to identify the TSS, provides more accurate annotations for the purpose of CRISPRi sgRNA selection than either NCBI (RefSeq) or Ensembl, which rely on alternative approaches to identify the TSS^32^. Based on our analysis of existing data^18,33^, and consistent with previous findings^19,32^, we identified the window of +25 to +75 nts downstream of the TSS as the optimal location for CRISPRi sgRNAs (Fig. 3a).

**Figure 3.**
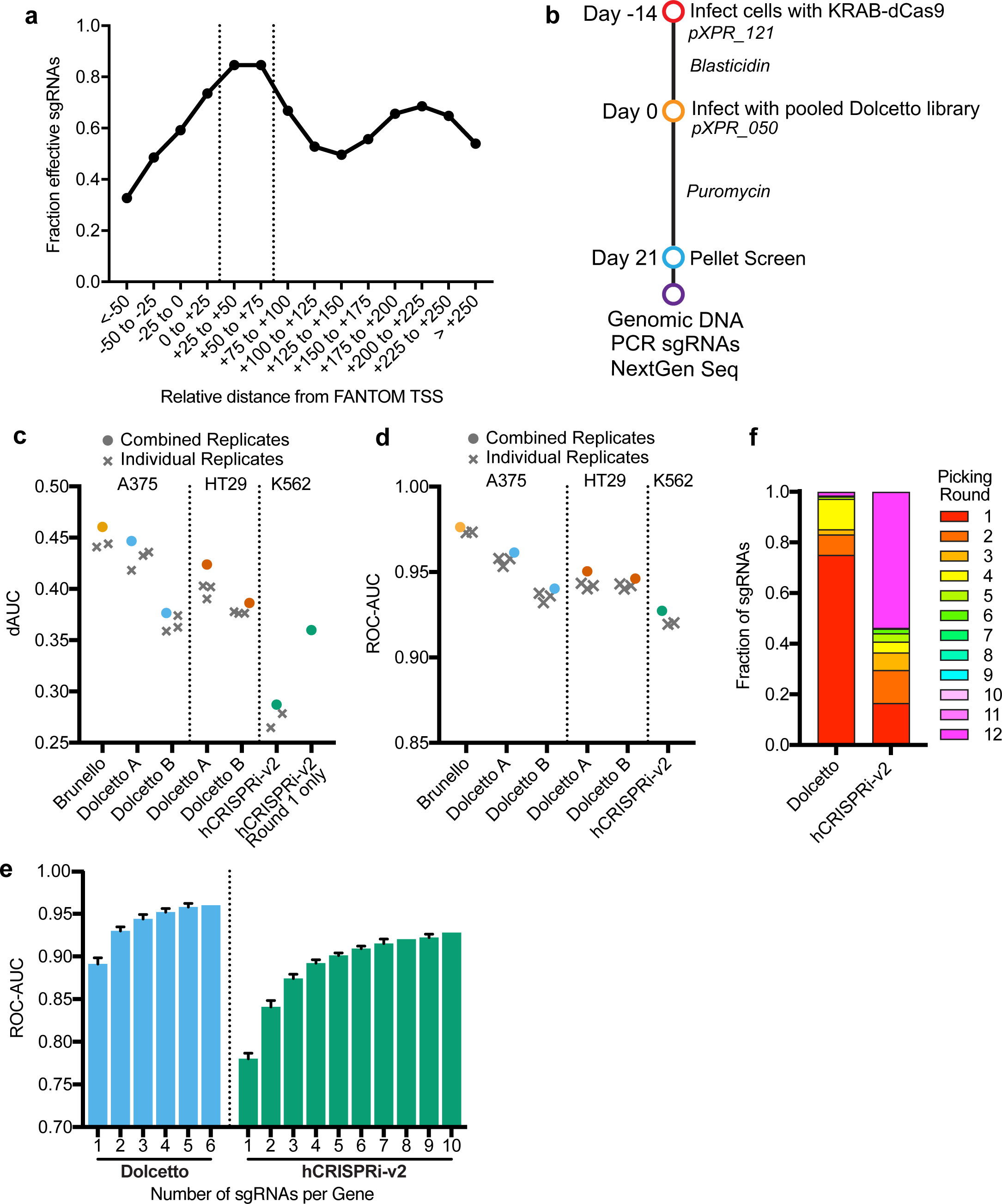
CRISPRi screening with Dolcetto. (a) Comparison of sgRNA activity as a function of distance from annotated transcription start site (TSS). Vertical dotted lines indicate the preferred targeting site. (b) Schematic of viability screens performed with Dolcetto in two cell lines. (c) dAUC comparison across CRISPRi libraries by cell line. dAUC values of individual replicates are plotted as X’s while dAUCs calculated using the average log2-fold change of replicates are plotted as circular points. For hCRISPR-v2 Round 1 only, the dAUC of the hCRISPR-v2 sgRNAs that would be picked in round 1 of our selection heuristic is plotted. Brunello dAUC values are plotted for comparison. (d) ROC-AUC comparison across CRISPRi libraries by cell line. ROC-AUC values of individual replicates are plotted as X’s while ROC-AUCs calculated using the average log2-fold change of replicates are plotted as circular points. Brunello ROC-AUC values are plotted for comparison. (e) Subsampling analysis, calculating the ROC-AUC of n sgRNAs per gene in Dolcetto and hCRISPRi-v2. Error bars represent different iterations of library sampling. (f) Fraction of sgRNAs selected in each selection round using the heuristics described in Supplementary Table 1 for Dolcetto and hCRISPRi-v2.

To design a CRISPRi library, we first selected from sgRNAs in this optimal window, and further ranked them by our previously-described sequence score, which has been shown to be effective for CRISPRi sgRNAs^16^, as well as their potential for off-target effects. In order to fulfill a quota of 6 sgRNAs per gene, we successively relaxed these three criteria (Supplementary Table 1). The resulting library, which we named Dolcetto, was divided into Sets A and B, with the former containing the top three sgRNAs selected by our heuristic. This library was cloned into a modified version of lentiGuide (pXPR_050). Here, we opted to use tracr-v2 for CRISPRi both because off-target effects are potentially less of a concern due to the limited window of activity and because previous CRISPRi studies have used a modified tracrRNA structure.

To assess the performance of Dolcetto, we generated A375 and HT29 cell lines stably expressing KRAB-dCas9, using a previously described lentiviral vector^34^, and performed negative selection screens in triplicate (Fig. 3b). We applied the dAUC (Fig. 3c) and ROC-AUC (Fig. 3d) metrics described above to compare performance. In both cell lines, Set A performed slightly better than Set B, indicating that the top three sgRNAs selected by our heuristic were indeed more likely to be active than the next three sgRNAs. We applied the same metrics to previously-published CRISPRi screens with the hCRISPRi-v2 library^19^ in K562 cells, and observed substantially better performance with Dolcetto (Fig. 3c, d). We next performed subsampling analysis to further compare libraries (combining Set A and Set B for Dolcetto). Even with only 3 sgRNAs per gene, Dolcetto outperformed hCRISPRi-v2 with a full 10 sgRNAs per gene (Fig. 3e), suggesting that the heuristics used to design this library were highly effective.

To further understand the relative importance of the various criteria in our design heuristic, we assigned the sgRNAs in hCRISPRi-v2 to the various selection rounds in our heuristic. We observed that a small fraction of hCRISPRi-v2 sgRNAs (17%) were in the first selection round, in contrast to Dolcetto, for which 75% of sgRNAs were selected in round one (Fig. 3f). We found that the dAUC for this filtered subset of the hCRISPRi-v2 library was higher than the original hCRISPRi-v2 library, with the dAUC improving from 0.29 for all guides to 0.36 for this subset (Fig. 3c). We therefore conclude that the differences in performance between hCRISPRi-v2 and Dolcetto can mostly be attributed to improved CRISPRi sgRNA design rather than differences across cell lines or experimental execution.

### Comparison of CRISPRko and CRISPRi

The dAUC and ROC-AUC metrics showed that Brunello and Dolcetto performed similarly in the discrimination between essential and non-essential genes. We next examined the data for signs of cutting-related toxicity, as has been previously been reported to be present with CRISPRko ^22-24^ but not with CRISPRi^19,34^. For this analysis, we compared the AUCs for sgRNAs targeting non-essential genes, which should cut the genome with Brunello but not Dolcetto, with non-targeting sgRNAs, which should not cut the genome with either library (Fig. 4a). Brunello exhibited a distinct cutting effect, as demonstrated by a lower AUC for non-targeting sgRNAs (AUC = 0.09) compared to sgRNAs targeting non-essential genes (0.41), as has been detailed previously^10^. Dolcetto, however, showed very little difference between non-essential (AUC = 0.41) and non-targeting sgRNAs (AUC = 0.45), indicating that CRISPRi mitigates cutting-related toxicity.

**Figure 4.**
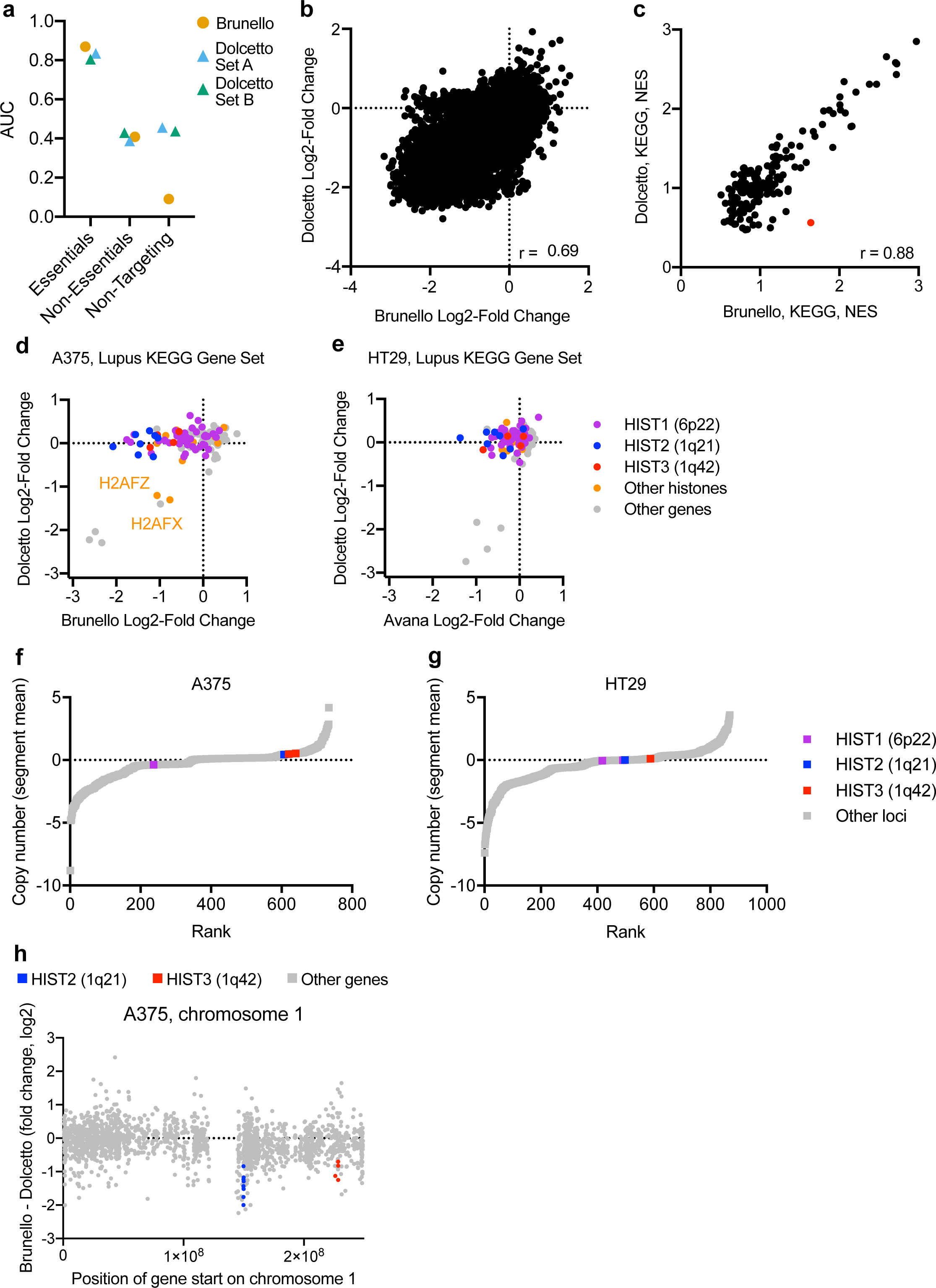
Comparisons across CRISPRko and CRISPRi. (a) Comparison of AUCs for essential, non-essential and non-targeting sgRNAs across CRISPRko and CRISPRi libraries screened in A375 cells. CRISPRko libraries are plotted as circular points and CRISPRi libraries are plotted as triangles. (b) Scatter plot comparing log2-fold change of genes in Brunello and Dolcetto (average of Sets A and B). Pearson correlation is reported. (c) Scatter plot comparing the GSEA normalized enrichment scores for KEGG genes sets for Brunello and Dolcetto. Pearson correlation is reported. (d) Scatter plot comparing log2-fold change of genes from the KEGG Systemic Lupus Erythematosus gene set for Brunello and Dolcetto in A375 cells. Histone genes are colored based on their chromosomal location. (e) Scatter plot comparing log2-fold change of genes from the KEGG Systemic Lupus Erythematosus gene set for Avana and Dolcetto in HT29 cells. Genes are colored as in (d). (f) Segmented copy number (segment mean; log2-fold change from average) of genetic loci in A375 cells. Probes that overlap histones in the Lupus gene set are labeled as in (d). (g) Segmented copy number (segment mean; log2-fold change from average) of genetic loci in HT29 cells. Probes are labeled as in (f). (h) Position of gene start along chromosome 1 compared to the difference in log2-fold change between Brunello and Dolcetto in A375 cells. Histone genes are colored as in (d).

Previous comparisons between CRISPRko, CRIPSRi, and RNAi suggested that the two technologies may identify different biological categories of essential genes^35,36^. To compare CRISPRi to CRISPRko, we first examined the gene-level correlation between Brunello and Dolcetto (Pearson r = 0.69; Fig. 4b). For any given gene, differences between essentiality between CRISPRko and CRISPRi may reasonably be attributed to differences in the efficacy of the individual sgRNAs rather than gene-intrinsic differences between knockout and knockdown. Therefore, to determine whether there were categories of genes that were systematically more depleted by one technology, we performed Gene Set Enrichment Analysis^37,38^ (GSEA) using the KEGG gene sets. We saw excellent correspondence between Brunello and Dolcetto (Pearson r = 0.88; Fig 4c), indicating that, at least in general, genes that manifest in a proliferation defect are equally assessed by both technologies when using optimized reagents.

Interestingly, one gene set, “Systemic Lupus Erythematosus,” was an outlier in this comparison, and thus we examined this in more detail. When we compared the performance of each individual gene in this set, we saw that numerous histone genes were essential when assessed by CRISPRko but not by CRISPRi in both A375 and HT29 cells (Fig. 4d, e). This observed difference between gene knockout and knockdown may represent a false positive finding with CRISPRko or a false negative result with CRISPRi. Most histone genes are found in three clusters on chromosomes 1 and 6^39^; a simple explanation would therefore be that these regions are copy number amplified and therefore show cutting toxicity with CRISPRko. However, neither region of chromosome 1 or 6 shows evidence of high copy number in A375 or HT29 cells (Fig. 4f-g). Additionally, several non-histone genes that are located near the histone clusters on chromosome 1 show comparable depletion with Brunello and Dolcetto in A375 cells, including both genes that are depleted by both technologies and depleted by neither (Fig. 4h). This further suggests that these regions are neither copy number amplified nor inaccessible to CRISPRi reagents. However, it is entirely possible that CRISPRi sgRNAs are ineffective at targeting histones; for example, there may be systematic biases in TSS annotation for these genes. Another unusual feature of histone genes is that most are only transcribed during S phase^40^. Intriguingly, in A375 cells, two notable exceptions to the differential depletion of histones were H2AFX and H2AFZ, two histone genes which are independently located on chromosomes 11 and 4, respectively, and unlike most histones, are transcribed in a replication-independent fashion. This raises the possibility that CRISPRi may be less effective at repressing gene expression during S phase. Finally, this observation may represent a true differential response to gene knockout and gene knockdown: perhaps even low levels of histone gene expression are sufficient for cell survival. These possibilities highlight that, despite the overall similarity of Brunello and Dolcetto, observed differences in response to the two technologies may shed light on interesting biological phenomena.

### Calabrese, a genome-wide CRISPRa library

The use of dCas9 also enables transcriptional activation (CRISPRa). Here, multiple strategies to recruit transcriptional machinery have proven effective^41^, including directly fusing activation domains to dCas9 (e.g. VP16), the use of the “Sun Tag” to recruit multiple copies of VP16^42^, or modifying the tracrRNA region of the sgRNA to include structured RNA domains such as MS2 that can then recruit additional transcription factors^20^. Based on previous studies optimizing the location of modifications to the tracrRNA^43^, we introduced two MS2 stem loops and two PP7 stem loops to create tracr-v14, a design that may allow higher-order combinations of domain recruitment (Fig. 5a). Since we encountered difficulties generating lentivirus of reasonable titer from the MS2-p65-HSF1 construct described in the original SAM system^20^, we opted to use a PP7-p65-HSF1 cassette, which was added to the sgRNA expression vector, to enable the use of only two vectors to conduct a CRISPRa screen.

**Figure 5.**
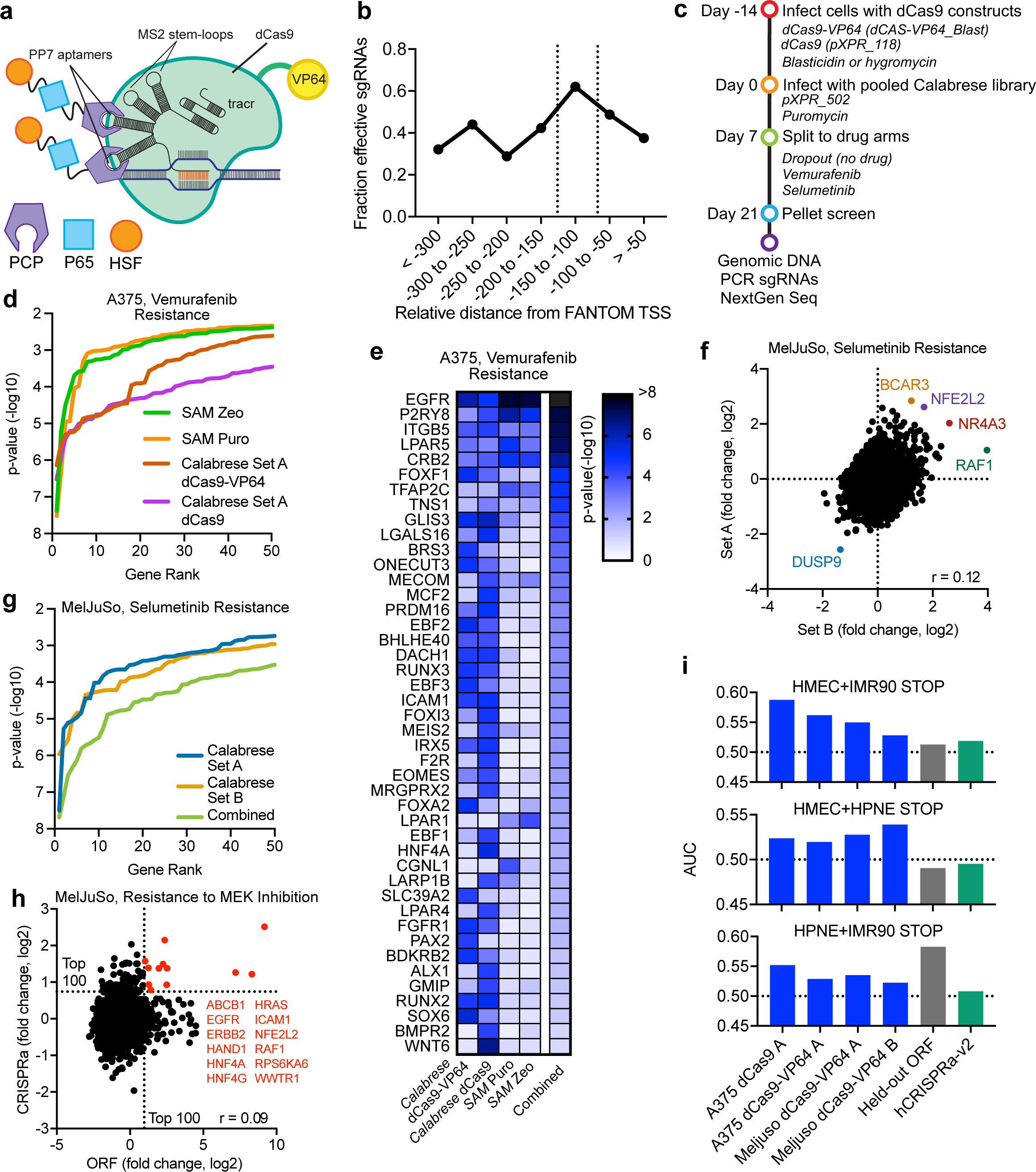
CRISPRa screening with Calabrese. (a) Schematic depicting PP7 and MS2 stem loop structure of tracr-v14. (b) Comparison of sgRNA activity as a function of distance from annotated transcription start site (TSS). Vertical dotted lines indicate the preferred targeting site. (c) Schematic of screens performed with Calabrese. (d) Line graph plotting the –log 10 p-value for the top 50 genes in vemurafenib resistance screens in A375 cells with the SAM and Calabrese libraries after averaging replicates. (e) Heat map of the most significant hits by p-value in CRISPRa vemurafenib resistance screens in A375 cells. The genes shown scored with p-value < 10^−4^ in any one of the four CRISPRa screens shown in (d) or in the combined analysis of all screens. (f) Scatter plot comparison of log2-fold change values for Calabrese Set A and Set B. Each dot represents the average of the 3 sgRNAs per gene. Pearson r correlation is reported. (g) Line graph plotting the –log 10 p-value for the top 50 genes in both sets of Calabrese and both sets combined for selumetinib resistance screens in MelJuSo cells. (h) Scatter plot comparison of log2-fold change for MEK inhibition screens in MelJuSo cells screened with an ORF library and trametinib and with the Calabrese library and selumetinib. Lines are drawn for the top 100 genes in each screen. (i) Area under the curve calculated for STOP genes in CRISPRa and held-out ORF screens. For each plot, a different ORF screen cell line (HMEC, IMR90, or HPNE) was held out, and the other two, indicated at the top of each graph, were used to generate a list of STOP genes.

As with CRISPRi, it is vital to consider location preference for CRISPRa sgRNAs for effective upregulation of genes. We used the same annotation database (FANTOM) for CRISPRa sgRNA design as we did for CRISPRi; however, for CRISPRa sgRNAs we targeted a window that was 150-75 nucleotides upstream of the TSS, based on re-analysis of previous data^18,33^ (Fig. 5b). We also used the same selection heuristics as for CRISPRi (Supplementary Table 2), successively relaxing location, on-target sequence score, and potential off-targets to select the six best sgRNAs for each gene, which were then divided into Set A (the top three sgRNAs) and Set B (the next three sgRNAs). This library, named Calabrese, was cloned into the pXPR_502 library vector, which contains the modified tracrRNA described above as well as the transcriptional activation domains p65-HSF1.

Unlike CRISPRko and CRISPRi, CRISPRa lacks an obvious gold standard gene set with which to assess screen performance in negative selection screens. Previously, the SAM system was screened for vemurafenib resistance in A375 cells^20^; this library contains 3 sgRNAs per gene and was screened in duplicate in two vectors, one of which conferred zeocin resistance and one puromycin resistance; the pairwise correlation for biological replicates with the same selection marker was 0.04 and 0.24, respectively. Low replicate correlation is generally expected in stringent positive selection screens, as the majority of sgRNAs do not confer resistance. To assess the performance of Calabrese, we performed a similar vemurafenib-resistance screen: we established A375 cells stably expressing dCas9-VP64^5,6,20^ and transduced with Calabrese Set A at low MOI (Fig. 5c). Additionally, we screened Calabrese Set A in A375 cells expressing dCas9 without the VP64 domain; here, recruitment of activation domains occurred solely via the PP7 stem loops in the library vector. Comparing the log2-fold change of vemurafenib-treated to untreated cells showed replicate correlation of 0.08 (dCas9-VP64) and 0.47 (dCas9). After collapsing sgRNAs to genes, we found that both Calabrese screens revealed substantially more hits at various p-value thresholds than either screen with the SAM library (Fig 5d), indicating better concordance of sgRNAs targeting the same gene. For example, at a p-value cut-off of 10^−4^, the Calabrese screens with dCas9-VP64 and dCas9 identified 17 and 27 genes, respectively, while the two SAM screens identified 4 and 5 genes each. We note that the original SAM library used RefSeq for annotating TSS and did not incorporate on-target selection rules, and the same sgRNAs were used in an updated version of the library^44^, which may partly explain its performance.

We next compared the performance of screens performed with dCas9-VP64 and dCas9 alone. Many of the same hit genes emerged from these screens, validating that dCas9 alone can be used for CRISPRa screens when activation domains are recruited by the library vector (Fig 5e). We then compared strong hits (p-value < 10^−4^) across all screens. One of the strongest hits with both the Calabrese and SAM libraries was EGFR, whose activation has previously been identified as a mediator of vemurafenib resistance^45,46^. Four other genes scored in all screens: P2RY8, ITGB5, LPAR5, and CRB2. LPAR5 belongs to the lysophosphatidic acid receptor family, which are G-protein coupled receptors that activate the PI3K-AKT signaling pathway^47^; other LPA receptors (LPAR1, LPAR4) also scored strongly in at least one screen. Additionally, a large number of genes scored only in Calabrese, including several groups of related genes, such as the transcription factors RUNX2 and RUNX3, as well as EBF1 and EBF2. Interestingly, in a genome-wide activation screen of lncRNAs using the SAM system, activation of the NR_109890 lncRNA was found to increase the expression of EBF1 and confer vemurafenib resistance^48^. When we combined data from both Calabrese and SAM library screens, several additional genes emerged as strong hits, including GLIS3 (Fig. 5e); knockdown of GLIS3 has been shown to enhance apoptosis, suggesting that GLIS3 activation could protect cells from apoptotic death^49^. In sum, the Calabrese library identify both previously-validated and novel genes that confer resistance to vemurafenib upon overexpression.

We next tested the performance of Calabrese in another cell line, MelJuSo, a BRAF-WT melanoma line first engineered to express dCas9-VP64. We performed a positive selection screen with selumetinib, a MEK inhibitor with both Set A and B in duplicate. Compared to the no-drug treatment control arm, pairwise Pearson correlation of log2-fold change values across biological replicates was 0.23 for Set A and 0.41 for Set B; after averaging together the three sgRNAs for each gene, the Pearson correlation comparing Set A to Set B was 0.12 (Fig. 5f). Each screen identified several of the same top hit genes that conferred selumetinib resistance upon activation, including the antioxidant responsive transcription factor NFE2L2 and the multidrug transporter ABCB1; conversely, activation of DUSP9, a MAPK phosphatase, sensitized cells to selumetinib. Sets A and B uncovered comparable numbers of genes at various p-value thresholds, and when Sets A and B were combined so that each gene was targeted with 6 sgRNAs, substantially more genes were uncovered at each statistical threshold (Fig. 5g). Thus, both sets of the Calabrese library were able to successfully identify genes conferring resistance to MAPK pathway inhibition.

### Comparison of CRISPRa to ORF overexpression

A previously-executed screen for resistance to MEK inhibition allowed us to compare the activity of a pooled open reading frame (ORF) overexpression library to CRISPRa^50,51^. This ORF screen was also performed in MelJuSo cells, although it used the MEK inhibitor trametinib rather than selumetinib. In that screen, across four replicates, the pairwise correlations of the log-fold-change values, relative to the no-drug treatment, ranged from 0.18 to 0.20. Considering the 11,960 genes shared in common across the two libraries, the correlation between the two screens at the gene level was modest (Fig. 5h, Pearson r = 0.09). However, common hits emerged. Of the top 100 resistance genes identified by each screen, 12 were found in both screens, including known MAPK oncogenes EGFR, RAF1, HRAS, and ERBB2, a statistically significant overlap (p-value 3×10^−11^, two-tailed Fisher’s exact test). Although we cannot rule out that some of the differences are due to the use of distinct MEK inhibitors, the data do not indicate that ORF and CRISPRa technologies are fully interchangeable, but rather uncover complementary hits, potentially due to higher false positive rates from both technologies relative to loss-of-function approaches.

A limitation of drug resistance screens for benchmarking library performance is that there is no orthogonal gold standard of what genes should score, in contrast to the core essential genes for loss-of-function technologies. We therefore sought to assess the effectiveness of Calabrese at modulating proliferation-related genes in the absence of drug treatment. A recent study using a large-scale ORF library of ~10,000 clones screened in 3 cancer cell lines identified growth inhibitory (STOP) and promoting (GO) genes^52^. Most STOP and GO genes were identified as cell-type specific, with only 103 and 3 genes, respectively, scoring in all 3 lines. Nevertheless, these data are a useful unbiased comparator for the performance of CRISPRa libraries in negative selection screens. Using the data provided in Sack et al., we generated 3 lists of STOP genes by holding out one cell line and requiring that the gene score in both of the remaining two cell lines. We then determined the AUC for the held-out ORF screen and the no-treatment arms of the Calabrese screens described above; we also analyzed a previously-described hCRISPRa-v2 screen performed in K562 cells^19^. Although the AUC values were modest in all cases, likely due to the cell-type specificity of most STOP genes, in all cases the AUC was >0.5 for the screens performed with Calabrese, indicating enrichment. Further, for two of the three lists of STOP genes, the Calabrese CRISPRa screens had a higher AUC than the held-out ORF screen, and in all cases outperformed the hCRISPRa-v2 library in this analysis (Fig. 5i). That sgRNAs activating growth inhibitory genes are preferentially depleted suggests that Calabrese is effective at modulating proliferation.

## DISCUSSION

Here we present three compact human genome-wide libraries for CRISPRko, CRISPRi, and CRISPRa and test their performance via 16 genome-wide screens conducted across 3 cell types. We demonstrate that these libraries outperform larger libraries on both the gene and sgRNA level, which we attribute to improved sgRNA design. Although we have not tested them here, we expect that the mouse versions of the CRISPRko, CRISPRa, and CRISPRi libraries, named Brie, Caprano, and Dolomiti, respectively, will also offer improved performance, as previous studies did not observe significant differences between sgRNA design criteria in mouse and human cells^9,12^. Indeed, a subset of the Brie library was screened *in vivo* in a model of tumor response to immunotherapy^53^, and another recent study successfully screened the Caprano library to identify anti-norovirus factors^54^.

Our results highlight the utility of modifying gene expression in multiple ways – up, down, and out – to fully probe gene function. We demonstrate that Brunello and Dolcetto, for CRISPRko and CRISPRi, respectively, are both highly effective loss-of-function libraries, with comparable depletion of essential genes. Additionally, we find a strong correlation between the gene sets depleted by these libraries, suggesting that optimized reagents may overcome some of the observed systematic differences between gene knockout by CRISPR and gene knockdown by RNAi^35^. One exception to the strong correlation between CRISPRko and CRISPRi was histone genes, which were preferentially depleted by CRISPRko; we speculate that this may reflect either a weakness of CRISPRi to repress genes that are transcribed during S phrase or a genuine biological response to low levels of histone gene expression. Additionally, although the technologies perform similarly overall, CRISPRko and CRISPRi each have weaknesses: in particular, CRISPRko leads to cutting-related toxicity at copy number amplified loci^24^, whereas CRISPRi can downregulate multiple genes at bidirectional promoters^34^. Each also has strengths; notably, CRISPRi offers the possibility of titrating the amount of gene expression to varying levels, which may prove useful for drug target discovery. Therefore, CRISPRko and CRISPRi screens represent valuable orthogonal approaches that can be leveraged to separate technical artifacts from true hits.

Upregulation of genes through CRISPRa likewise represents a complementary method for pooled screening. In contrast to gene knockout or knockdown screens, CRISPRa screens can reveal the function of genes with a low baseline level of expression and uncover pathways that are more effectively modulated by gene activation. Additionally, CRISPRa is a powerful approach for strong positive selection screens, for which it is often easiest to separate signal from noise for resistance hits; CRISPRa screens may therefore strongly identify genes that may score weakly on the sensitization side in knockout screens but represent a true biological hit. CRISPRa screens are also complementary to ORF screens. When we compared CRISPRa and ORF screens for resistance to MEK inhibitors, we found that the technologies shared a number of common top hits, but also identified numerous unique ones. Both technologies have sources of false negatives and positives that may explain this difference. Sources of false negative findings for ORF technology include overexpression of a splice isoform that is functionally inactive in the assay of interest; for CRISPRa, false negatives may arise when poor sgRNA design leads to failure to overexpress the target gene or when local chromatin environment renders a gene inaccessible to Cas9. False positives can occur in ORF screens when an ORF is overexpressed to a level not ever achieved by the cell endogenously and, in CRISPRa, when multiple genes are upregulated at bidirectional promoters. Therefore, both ORF and CRISPRa screens are valuable and complementary elements of the pooled screening toolbox.

Another factor in screen performance is the design of the tracrRNA. Here, we demonstrate that a modified tracrRNA, tracr-v2, performs comparably to the original tracrRNA in a genome-wide screen but shows increased on-target activity and increased off-target effects when screened with a tiling library. We therefore conclude that either the original tracrRNA or tracr-v2 may be used for screening, although tracr-v2 may be preferable in cases (such as CRISPRi) where off-target effects are less of a concern. Additionally, we performed CRISPRa screens with a dCas9 in which activation domains were only recruited by PP7 stem loops in the tracrRNA. The strong performance of this approach indicates that it may be possible to use a single dCas9-expressing cell line for both CRISPRa and CRISPRi screens when modulatory domains are recruited by the library vector. This approach would be especially beneficial for screens in primary cells, where establishment of a Cas9-expressing cell line can be challenging. Additionally, previous reports have demonstrated that a truncated sgRNA can be used to bind Cas9 without inducing cleavage activity^55,56^. Therefore, it may be possible to establish a single Cas9 cell line and screen a library that includes full-length sgRNAs for CRISPRko, truncated sgRNAs in a vector that recruits inhibitory domains for CRISPRi, and truncated sgRNAs in a vector that recruits activating domains for CRISPRa.

Although CRISPR-Cas9 technology has greatly expanded the capacity to conduct large-scale, unbiased screens to characterize gene function, designing screenable model systems remains a challenge for many disease areas; often, primary cells or *in vivo* models best capture relevant disease biology, but are limited in cell number and therefore library size. Moreover, once a high-quality, screenable model system has been developed, it is often advantageous to do many screens, as different technologies can each uncover distinct hits. The highly compact CRISPRko, CRISPRi, and CRISPRa libraries introduced here will enable genome-scale interrogation of gene function in diverse model systems and assist in expanding the scope of pooled screening beyond easily-cultured cell lines.

## ACKNOWLEDGMENTS

We are grateful to be supported by excellent software engineers: Matthew Greene, Doug Alan, Mark Tomko, Adam Brown, and Tom Green; as well as administrative support: Olivia Baré and Samantha Amaral (Genetic Perturbation Platform, Broad Institute). We thank Tamara Mason and the Walk-Up Sequencing Team for high quality and rapid Illumina sequencing (Genomics Platform, Broad Institute); Tikvak Hayes and Cory Johannessen for sharing the trametinib-resistance ORF screening data prior to publication, and Mahmoud Ghandi for helpful discussions on the interpretation of copy number data (Cancer Program, Broad Institute). We also thank Laura Sack and Steve Elledge (Harvard Medical School) for helpful guidance on the interpretation of their ORF screening data. This work was supported in part by the Functional Genomics Consortium.

## METHODS

### Vectors

The following vectors were used in this study and are available to the academic community on Addgene:

pLX_311-Cas9: SV40 promoter expresses blasticidin resistance; EF1a promoter expresses SpCas9 (Addgene 96924).

dCAS-VP64_Blast (pXPR_109): EF1a promoter expresses dSpCas9-VP64 and 2A site provides blasticidin resistance (Addgene 61425).

pXPR_118: EF1a promoter expresses dSpCas9 and 2A site provides hygromycin resistance (Addgene, in process).

pXPR_121: SV40 promoter expresses blasticidin resistance; EF1a promoter expresses KRAB-dSpCas9 (Addgene 96918).

lentiGuide-Puro (pXPR_003): EF1a promoter expresses puromycin resistance; U6 promoter expresses customizable sgRNA element (Addgene 52963).

lentiCRISPRv2 (pXPR_023): EF1a promoter expresses SpCas9 and 2A site provides puromycin resistance; U6 promoter expresses customizable sgRNA element (Addgene 52961).

pXPR_050: EF1a promoter expresses puromycin resistance; U6 promoter expresses customizable sgRNA element with tracr-v2 (Addgene 96925).

pXPR_502: PGK promoter expresses PP7 tethered to P65-HSF1 and 2A site provides puromycin resistance; U6 promoter expresses customizable sgRNA element with tracr-v14, containing 2 MS2 and 2 PP7 stem loops (Addgene 96923)

Brunello library: Addgene 73178 (in lentiGuide-Puro) and 73179 (in lentiCRIPSRv2)

Calabrese library: Addgene 92379 (Set A), 92380 (Set B)

Dolcetto library: Addgene 92385 (Set A), 92386 (Set B)

### CRISPRi and CRISPRa library design

We used the FANTOM5 dataset to define transcription start sites (TSS) for the genes defined as protein-coding in the human genome by RefSeq. For each gene with TSS information in FANTOM, we picked the TSS with the highest-ranked peak of all the CAGE peaks. For the remaining genes, we used the TSS information from Ensembl or NCBI.

For each gene, we first designed all possible sgRNAs with NGG PAM in the window of −300 to +300 relative to the TSS. We then calculated the Rule Set 2 scores for these sgRNAs and annotated each sgRNA with the number of perfectly matched off-target sites in the human genome.

We then picked sgRNAs based on their position relative to the TSS, number of off-target matches and on-target score to fill a quota of six sgRNAs per gene. (Supplementary Table 1 for CRISPRi and Supplementary Table 2 for CRISPRa) One or more of these three criteria were relaxed in each picking round until the quota was filled. A standard set of 992 non-targeting controls was added to each library. We then divided each library into Set A and Set B with the former having the top 3 sgRNAs and the latter having the next 3 sgRNAs per gene.

### Library production

Libraries were prepared as described previously^16^. Briefly, oligonucleotide pools were synthesized at CustomArray. BsmBI recognition sites were appended to each sgRNA sequence along with the appropriate overhang sequences (underlined) for cloning into the sgRNA expression plasmids, as well as primer sites to allow differential amplification of subsets from the same synthesis pool. The final oligonucleotide sequence was thus: 5’-[Forward Primer]CGTCTCACACCG[sgRNA, 20 ntlGTTTCGAGACG[Reverse Primer].

Primers were used to amplify individual subpools using 25 μL 2x NEBnext PCR master mix (New England Biolabs), 2 μL of oligonucleotide pool (~40 ng), 5 μL of primer mix at a final concentration of 0.5 μM, and 18 μL water. PCR cycling conditions: 30 seconds at 98°C, 30 seconds at 53°C, 30 seconds at 72°C, for 24 cycles. In cases where a library was divided into subsets unique primers could be used for amplification:

Primer Set; Forward Primer, 5’ - 3’; Reverse Primer, 5’ - 3’

1. AGGCACTTGCTCGTACGACG; ATGTGGGCCCGGCACCTTAA
2. GTGTAACCCGTAGGGCACCT; GTCGAGAGCAGTCCTTCGAC
3. CAGCGCCAATGGGCTTTCGA; AGCCGCTTAAGAGCCTGTCG
4. CTACAGGTACCGGTCCTGAG; GTACCTAGCGTGACGATCCG
5. CATGTTGCCCTGAGGCACAG; CCGTTAGGTCCCGAAAGGCT
6. GGTCGTCGCATCACAATGCG; TCTCGAGCGCCAATGTGACG

The resulting amplicons were PCR-purified (Qiagen) and cloned into the library vector via Golden Gate cloning with Esp3I (Fisher Scientific) and T7 ligase (Epizyme); the library vector was pre-digested with BsmBI (New England Biolabs). The ligation product was isopropanol precipitated and electroporated into Stbl4 electrocompetent cells (Life Technologies) and grown at 30°C for 16 hours on agar with 100 μg/mL carbenicillin. Colonies were scraped and plasmid DNA (pDNA) was prepared (HiSpeed Plasmid Maxi, Qiagen). To confirm library representation and distribution, the pDNA was sequenced by Illumina.

### Virus production

Virus production was carried out as described previously^16^. For individual virus production, the following procedure was used: 24 h before transfection, HEK293T cells were seeded in 6-well dishes at a density of 1.5 × 10^6^ cells per well in 2 mL of DMEM + 10% FBS. Transfection was performed using TransIT-LT1 (Mirus) transfection reagent according to the manufacturer’s protocol. Briefly, one solution of Opti-MEM (Corning, 66.25 μL) and LT1 (8.75 μL) was combined with a DNA mixture of the packaging plasmid pCMV_VSVG (Addgene 8454, 250 ng), psPAX2 (Addgene 12260, 1,250 ng), and the transfer vector (e.g., pLentiGuide, 1,250 ng). The solutions were incubated at room temperature for 20-30 min, during which time media was changed on the HEK293T cells. After this incubation, the transfection mixture was added dropwise to the surface of the HEK293T cells, and the plates were centrifuged at 1,000g for 30 min at room temperature. Following centrifugation, plates were transferred to a 37 °C incubator for 6-8 h, after which the media was removed and replaced with DMEM + 10% FBS media supplemented with 1% BSA.

A larger-scale procedure was used for pooled library production. 24 h before transfection, 18 × 10^6^ HEK293T cells were seeded in a 175 cm^2^ tissue culture flask and the transfection was performed as described above using 6 mL of Opti-MEM, 305 μL of LT1, and a DNA mixture of pCMV_VSVG (5 μg), psPAX2 (50 μg), and 40 μg of the transfer vector. Flasks were transferred to a 37 °C incubator for 6-8 h; after this, the media was aspirated and replaced with BSA-supplemented media. Virus was harvested 36 h after this media change.

### Cell culture

A375, HT29, and MelJuSo cells were obtained from the Cancer Cell Line Encyclopedia. HEK293Ts were obtained from ATCC (CRL-3216).

All cell lines were routinely tested for mycoplasma contamination and were maintained without antibiotics except during screens, when media was supplemented with 1% penicillin/streptomycin. Cell lines were kept in a 37 °C humidity-controlled incubator with 5.0% CO2 and were maintained in exponential phase growth by passaging every 2-3 days.

For each cell line, the following media and doses of polybrene, puromycin, blasticidin, and hygromycin, respectively, were used:

A375: RPMI + 10% FBS; 1 μg/mL (0.5 μg/mL for no-spin transductions); 1 μg/mL; 5 μg/mL; 50 μg/mL

HT29: DMEM + 10% FBS; 1 μg/mL; 2 μg/mL; 5 μg/mL; N/A MelJuSo: RPMI + 10% FBS; 4 μg/mL; 1 μg/mL; 2 μg/mL; N/A

Vemurafenib (S1267) and selumetinib (S1008) were obtained from Selleckchem and screened at doses of 2 μM and 1.5 μM, respectively. 6-thioguanine (A4882) was obtained from Sigma-Aldrich and screened at a dose of 5 μg/mL.

### Screening

#### Determination of lentiviral titer

To determine lentiviral titer for spin transductions, cell lines were transduced in 12-well plates with a range of virus volumes (e.g. 0, 150, 300, 500, and 800 μL virus) with 3.0 × 10^6^ cells per well in the presence of polybrene. The plates were centrifuged at 640 × g for 2 h and were then transferred to a 37 °C incubator for 4-6 h. Each well was then trypsinized, and an equal number of cells seeded into each of two wells of a 6-well dish. Two days post-transduction, puromycin was added to one well out of the pair. After 5 days, both wells were counted for viability. A viral dose resulting in 30-50% transduction efficiency, corresponding to an MOI of ~0.35-0.70, was used for subsequent library screening.

To determine lentiviral titer for no-spin transductions, cell lines were seeded in 6-well plates in the presence of polybrene (0.5 μg/mL) and virus at a range of volumes (e.g. 0, 50, 100, 200, 400, 600 μL virus), with two wells per virus volume. 16-18 h after seeding, virus-containing media was replaced with fresh media. Two days post-transduction, puromycin was added to one well out of the pair. After 5 days, both wells were counted for viability. A viral dose resulting in 30-50% transduction efficiency, corresponding to an MOI of ~0.35-0.70, was used for subsequent library screening.

#### CRISPRko and CRISPRi screens

Cas9 or dCas9 derivatives were made by transducing with the lentiviral vector pLX_311-Cas9, which expresses blasticidin resistance from the SV40 promoter and Cas9 from the EF1α promoter, or pXPR_121 which expresses blasticidin resistance from the SV40 promoter and KRAB-dCas9 from the EF1a promoter, respectively.

Prior to screening-scale transduction, Cas9 and KRAB-dCas9-expressing cell lines were selected with blasticidin then transduced in two or three biological replicates at a low MOI (~0.5). Transductions were performed with enough cells to achieve a representation of at least 500 cells per sgRNA per replicate, taking into account a 30-50% transduction efficiency. Throughout the screen, cells were split at a density to maintain a representation of at least 500 cells per sgRNA, and cell counts were taken at each passage to monitor growth. Puromycin selection was added 2 days post-transduction and was maintained for 5-7 days. 3 weeks posttransfection, cells were pelleted by centrifugation, resuspended in PBS, and frozen promptly for genomic DNA isolation.

#### CRISPRa screens

Cell lines expressing dCas9-VP64 and dCas9 were made by transducing cells with the lentiviral vector pXPR_109, which expresses blasticidin resistance from a 2A site and dCas9-VP64 from the EF1α promoter, or pXPR_118, which expresses hygromycin resistance from a 2A site and dCas9 from the EF1α promoter, respectively. Prior to screening-scale transduction, dCas9-VP64 and dCas9 cell lines were selected with blasticidin and hygromycin, respectively.

For the dCas9-VP64 screens, cell lines expressing dCas9-VP64 were transduced with the Calabrese library in two biological replicates at a low MOI (~0.5). Transductions were performed with enough cells to achieve a representation of at least 500 cells per sgRNA per replicate, taking into account a 30-50% transduction efficiency. Throughout the screen, cells were split at a density to maintain a representation of at least 500 cells per sgRNA, and cell counts were taken at each passage to monitor growth. Puromycin selection was added 2 days posttransduction and was maintained for 5-7 days. After puromycin selection was complete, each replicate was split to no drug and drug treatment arms, each at a representation of at least 500 cells per sgRNA. A375 screens were performed with 2 μM vemurafenib; MelJuSo screens were performed with 1.5 μM selumetinib. 14 days after the initiation of drug treatment, cells were pelleted by centrifugation, resuspended in PBS, and frozen promptly for genomic DNA isolation.

For the dCas9-only screens, A375 cells expressing dCas9 were transduced with the Calabrese library in two biological replicates at a low MOI (~0.5) via a low-representation, no-spin method. Transductions were performed to achieve a representation of at least 300 sgRNAs per replicate, taking into account a 30-50% transduction efficiency. Briefly, cells per were seeded into T175 flasks in a total volume of 20 mL of media containing virus and polybrene at 0.5 μg/mL. Flasks were then transferred to an incubator overnight. 16-18 h after seeding, the virus-containing media was replaced with fresh media and cells were expanded to the average representation of at least 500 transduced cells per sgRNA. Puromycin selection was added 2 days post-transduction and was maintained for 5-7 days. After puromycin selection was complete, each replicate was split to no drug and drug treatment arms, each at a representation of at least 500 cells per sgRNA. 14 days after the initiation of drug treatment, cells were pelleted by centrifugation, resuspended in PBS, and frozen promptly for genomic DNA isolation.

### Genomic DNA preparation and sequencing

Genomic DNA (gDNA) was isolated using the QIAamp DNA Blood Maxi (3e7 - 1e8 cells), Midi (5e6 - 3e7 cells), or Mini (<5e6 cells) Kits (Qiagen) as per the manufacturer’s instructions. The gDNA concentrations were quantitated by UV Spectroscopy (Nanodrop). For PCR amplification, gDNA was divided into 100 μL reactions such that each well had at most 10 μg of gDNA. Per 96 well plate, a master mix consisted of 75 μL ExTaq DNA Polymerase (Clontech), 1000 μL of 10x Ex Taq buffer, 800 μL of dNTP provided with the enzyme, 50 μL of P5 stagger primer mix (stock at 100 μM concentration), and 2075 μL water. Each well consisted of 50 μL gDNA plus water, 40 μL PCR master mix, and 10 μL of a uniquely barcoded P7 primer (stock at 5 μM concentration). PCR cycling conditions: an initial 1 minute at 95°C; followed by 30 seconds at 94°C, 30 seconds at 52.5°C, 30 seconds at 72°C, for 28 cycles; and a final 10 minute extension at 72°C. P5/P7 primers were synthesized at Integrated DNA Technologies (IDT). PCR products were purified with Agencourt AMPure XP SPRI beads according to manufacturer’s instructions (Beckman Coulter, A63880). Samples were sequenced on a HiSeq2000 (Illumina), loaded with a 5% spike-in of PhiX DNA.

Reads were counted by first searching for the CACCG sequence in the primary read file that appears in the vector 5’ to all sgRNA inserts. The next 20 nts are the sgRNA insert, which was then mapped to a reference file of all possible sgRNAs present in the library. The read was then assigned to a condition (e.g. a well on the PCR plate) on the basis of the 8nt barcode included in the P7 primer.

### Screen analysis

Following deconvolution, the resulting matrix of read counts was first normalized to a reads per million within each condition by the following formula: read per sgRNA / total reads per condition × 10^6^. Reads per million was then log2-transformed by first adding one to all values, which is necessary in order to take the log of sgRNAs with zero reads. For each sgRNA, the log2-fold-change from plasmid DNA (pDNA) was then calculated. All reported log2-fold-changes for dropout screens are relative to pDNA; for positive selection screens with small molecules, the log2-fold-change relative to the dropout arm (i.e. no drug arm) was also calculated.

To determine p-values for genes to evaluate positive selection screens, we calculated the hypergeometric distribution without replacement based on the rank order of the log-fold-change of the perturbations; this is equivalent to a one-sided Fisher’s exact test. Non-targeting control sgRNAs were randomly grouped into dummy genes of the same set size as the library under consideration (e.g. 4 control sgRNAs for Brunello, 3 for Calabrese Set A, 6 for Calabrese Set A and B combined). The SAM screening data described previously^20^ was reanalyzed using this approach.

### Gene set enrichment analysis and analysis of lupus gene set

Gene set enrichment analysis (GSEA) was performed using the KEGG gene sets. In order to further investigate the genes in the Systemic Lupus Erythematosus gene set, we compared the depletion of genes in the gene set by CRISPRi (Dolcetto) to CRISPRko; for A375 cells, we used CRISPRko data from screens with Brunello, whereas for HT29 cells, we used CRISPRko data from Project Achilles, in which the Avana library was screened with 4 sgRNAs per gene.

To analyze copy number amplification in A375 and HT29 cells, we obtained segmented copy number data from Project Achilles^23^ (https://portals.broadinstitute.org/achilles). Gene position and karyotype band data were obtained from Ensembl.

### Analysis of STOP & GO genes

Using Supplementary Table 2B from the report by Sack et al., we identified STOP genes by requiring first that the ORF was screened in all three cell lines (no ‘n/a’ values). We then held out each cell line at a time, and required that the ORF clone have a negative log2 value (e.g. Column E) and a p-value <10^−4^ (e.g. Column F) in both other cell lines. Using the resulting list of genes as a reference of essential genes, we then calculated the area-under-the-curve for CRISPRa libraries and the held-out cell line from the ORF dataset.

### Code availability

All custom Python code used for analysis is available on GitHub: https://github.com/mhegde.

## FIGURE LEGENDS

**Supplementary Table 1**. CRISPRi library design picking criteria. CRISPRi sgRNAs were picked in 12 rounds binned by position relative to TSS, off-target criteria and on-target criteria.

**Supplementary Table 2**. CRISPRa library design picking criteria. CRISPRa sgRNAs were picked in 12 rounds binned by position relative to TSS, off-target criteria and on-target criteria.

CRISPRi library design picking criteria

CRISPRi library design picking criteria
| **Picking Round** | **Position relative to TSS** | **Off-target criteria** | **On-target criteria - Rule Set2 score** |
| --- | --- | --- | --- |
| 1 | +25 to +75 | $\leq 1$ perfect match | $\geq 0.2$ |
| 2 | 0 to +100 | $\leq 1$ perfect match | $\geq 0.2$ |
| 3 | +175 to +250 | $\leq 1$ perfect match | $\geq 0.2$ |
| 4 | +25 to +75 | $\leq 5$ perfect match | $\geq 0.2$ |
| 5 | 0 to +100 | $\leq 5$ perfect match | $\geq 0.2$ |
| 6 | +175 to +250 | $\leq 5$ perfect match | $\geq 0.2$ |
| 7 | +25 to +75 | $\leq 5$ perfect match | $> 0$ |
| 8 | 0 to +100 | $\leq 5$ perfect match | $> 0$ |
| 9 | +175 to +250 | $\leq 5$ perfect match | $> 0$ |
| 10 | -50 to +300 | $\leq 1$ perfect match | $> 0$ |
| 11 | -50 to +300 | $\leq 1$ perfect match | $> 0$ |
| 12 | -300 to +300 | Any | Any |

CRISPRa library design picking criteria
| **Picking Round** | **Position relative to TSS** | **Off-target criteria** | **On-target criteria - Rule Set2 score** |
| --- | --- | --- | --- |
| 1 | -150 to -75 | $\leq 1$ perfect match | $\geq 0.2$ |
| 2 | -200 to -25 | $\leq 1$ perfect match | $\geq 0.2$ |
| 3 | -250 to 0 | $\leq 1$ perfect match | $\geq 0.2$ |
| 4 | -150 to -75 | $\leq 5$ perfect match | $\geq 0.2$ |
| 5 | -200 to -25 | $\leq 5$ perfect match | $\geq 0.2$ |
| 6 | -250 to 0 | $\leq 5$ perfect match | $\geq 0.2$ |
| 7 | -150 to -75 | $\leq 5$ perfect match | $> 0$ |
| 8 | -200 to -25 | $\leq 5$ perfect match | $> 0$ |
| 9 | -250 to 0 | $\leq 5$ perfect match | $> 0$ |
| 10 | -250 to 0 | $\leq 1$ perfect match | $> 0$ |
| 11 | -300 to 0 | $\leq 1$ perfect match | $> 0$ |
| 12 | -300 to +300 | Any | Any |

## REFERENCES

1. Doench, J. G. Am I ready for CRISPR? A user’s guide to genetic screens. Nat. Rev. Genet. 19, 67–80 (2018).

2. Shalem, O. et al. Genome-scale CRISPR-Cas9 knockout screening in human cells. Science 343, 84–87 (2014).

3. Zalatan, J. G. et al. Engineering complex synthetic transcriptional programs with CRISPR RNA scaffolds. Cell 160, 339–350 (2015).

4. Gilbert, L. A. et al. CRISPR-mediated modular RNA-guided regulation of transcription in eukaryotes. Cell 154, 442–451 (2013).

5. Perez-Pinera, P. et al. RNA-guided gene activation by CRISPR-Cas9-based transcription factors. Nat. Methods 10, 973–976 (2013).

6. Maeder, M. L. et al. CRISPR RNA-guided activation of endogenous human genes. Nat. Methods 10, 977–979 (2013).

7. Gao, X. et al. Comparison of TALE designer transcription factors and the CRISPR/dCas9 in regulation of gene expression by targeting enhancers. Nucleic Acids Res. 42, e155 (2014).

8. Qi, L. S. et al. Repurposing CRISPR as an RNA-guided platform for sequence-specific control of gene expression. Cell 152, 1173–1183 (2013).

9. Doench, J. G. et al. Rational design of highly active sgRNAs for CRISPR-Cas9-mediated gene inactivation. Nat. Biotechnol. 32, 1262–1267 (2014).

10. Hart, T. et al. Evaluation and Design of Genome-Wide CRISPR/SpCas9 Knockout Screens. G3 7, 2719–2727 (2017).

11. Ong, S. H., Li, Y., Koike-Yusa, H. & Yusa, K. Optimised metrics for CRISPR-KO screens with second-generation gRNA libraries. Sci. Rep. 7, 7384 (2017).

12. Haeussler, M. et al. Evaluation of off-target and on-target scoring algorithms and integration into the guide RNA selection tool CRISPOR. Genome Biol. 17, 148 (2016).

13. Wang, T., Wei, J. J., Sabatini, D. M. & Lander, E. S. Genetic screens in human cells using the CRISPR-Cas9 system. Science 343, 80–84 (2014).

14. Tzelepis, K. et al. A CRISPR Dropout Screen Identifies Genetic Vulnerabilities and Therapeutic Targets in Acute Myeloid Leukemia. Cell Rep. 17, 1193–1205 (2016).

15. Hart, T. et al. High-Resolution CRISPR Screens Reveal Fitness Genes and Genotype-Specific Cancer Liabilities. Cell 163, 1515–1526 (2015).

16. Doench, J. G. et al. Optimized sgRNA design to maximize activity and minimize off-target effects of CRISPR-Cas9. Nat. Biotechnol. 34, 184–191 (2016).

17. Wang, T. et al. Gene Essentiality Profiling Reveals Gene Networks and Synthetic Lethal Interactions with Oncogenic Ras. Cell 168, 890–903.e15 (2017).

18. Gilbert, L. A. et al. Genome-Scale CRISPR-Mediated Control of Gene Repression and Activation. Cell 159, 647–661 (2014).

19. Horlbeck, M. A. et al. Compact and highly active next-generation libraries for CRISPR-mediated gene repression and activation. Elife 5, (2016).

20. Konermann, S. et al. Genome-scale transcriptional activation by an engineered CRISPR-Cas9 complex. Nature 517, 583–588 (2015).

21. Hart, T., Brown, K. R., Sircoulomb, F., Rottapel, R. & Moffat, J. Measuring error rates in genomic perturbation screens: gold standards for human functional genomics. Mol. Syst. Biol. 10, 733 (2014).

22. Munoz, D. M. et al. CRISPR Screens Provide a Comprehensive Assessment of Cancer Vulnerabilities but Generate False-Positive Hits for Highly Amplified Genomic Regions. Cancer Discov. 6, 900–913 (2016).

23. Aguirre, A. J. et al. Genomic Copy Number Dictates a Gene-Independent Cell Response to CRISPR/Cas9 Targeting. Cancer Discov. 6, 914–929 (2016).

24. Wang, T. et al. Identification and characterization of essential genes in the human genome. Science 350, 1096–1101 (2015).

25. Tsherniak, A. et al. Defining a Cancer Dependency Map. Cell 170, 564–576.e16 (2017).

26. Meyers, R. M. et al. Computational correction of copy number effect improves specificity of CRISPR-Cas9 essentiality screens in cancer cells. Nat. Genet. (2017). doi:10.1038/ng.3984

27. Chen, B. et al. Dynamic imaging of genomic loci in living human cells by an optimized CRISPR/Cas system. Cell 155, 1479–1491 (2013).

28. Dang, Y. et al. Optimizing sgRNA structure to improve CRISPR-Cas9 knockout efficiency. Genome Biol. 16, 280 (2015).

29. Cross, B. C. S. et al. Increasing the performance of pooled CRISPR-Cas9 drop-out screening. Sci. Rep. 6, 31782 (2016).

30. Najm, F. J. et al. Orthologous CRISPR-Cas9 enzymes for combinatorial genetic screens. Nat. Biotechnol. 36, 179–189 (2018).

31. FANTOM Consortium and the RIKEN PMI and CLST (DGT) et al. A promoter-level mammalian expression atlas. Nature 507, 462–470 (2014).

32. Radzisheuskaya, A., Shlyueva, D., Müller, I. & Helin, K. Optimizing sgRNA position markedly improves the efficiency of CRISPR/dCas9-mediated transcriptional repression. Nucleic Acids Res. 44, e141 (2016).

33. Wong, N., Liu, W. & Wang, X. WU-CRISPR: characteristics of functional guide RNAs for the CRISPR/Cas9 system. Genome Biol. 16, 218 (2015).

34. Rosenbluh, J. et al. Complementary information derived from CRISPR Cas9 mediated gene deletion and suppression. Nat. Commun. 8, 15403 (2017).

35. Morgens, D. W., Deans, R. M., Li, A. & Bassik, M. C. Systematic comparison of CRISPR/Cas9 and RNAi screens for essential genes. Nat. Biotechnol. 34, 634–636 (2016).

36. Evers, B. et al. CRISPR knockout screening outperforms shRNA and CRISPRi in identifying essential genes. Nat. Biotechnol. 34, 631–633 (2016).

37. Mootha, V. K. et al. PGC-1alpha-responsive genes involved in oxidative phosphorylation are coordinately downregulated in human diabetes. Nat. Genet. 34, 267–273 (2003).

38. Subramanian, A. et al. Gene set enrichment analysis: a knowledge-based approach for interpreting genome-wide expression profiles. Proc. Natl. Acad. Sci. U. S. A. 102, 15545–15550 (2005).

39. Marzluff, W. F., Gongidi, P., Woods, K. R., Jin, J. & Maltais, L. J. The human and mouse replication-dependent histone genes. Genomics 80, 487–498 (2002).

40. Nishida, H., Suzuki, T., Ookawa, H., Tomaru, Y. & Hayashizaki, Y. Comparative analysis of expression of histone H2a genes in mouse. BMC Genomics 6, 108 (2005).

41. Chavez, A. et al. Comparison of Cas9 activators in multiple species. Nat. Methods 13, 563–567 (2016).

42. Tanenbaum, M. E., Gilbert, L. A., Qi, L. S., Weissman, J. S. & Vale, R. D. A protein-tagging system for signal amplification in gene expression and fluorescence imaging. Cell 159, 635–646 (2014).

43. Shechner, D. M., Hacisuleyman, E., Younger, S. T. & Rinn, J. L. Multiplexable, locus-specific targeting of long RNAs with CRISPR-Display. Nat. Methods 12, 664–670 (2015).

44. Joung, J. et al. Genome-scale CRISPR-Cas9 knockout and transcriptional activation screening. Nat. Protoc. 12, 828–863 (2017).

45. Corcoran, R. B. et al. EGFR-mediated re-activation of MAPK signaling contributes to insensitivity of BRAF mutant colorectal cancers to RAF inhibition with vemurafenib. Cancer Discov. 2, 227–235 (2012).

46. Prahallad, A. et al. Unresponsiveness of colon cancer to BRAF(V600E) inhibition through feedback activation of EGFR. Nature 483, 100–103 (2012).

47. Riaz, A., Huang, Y. & Johansson, S. G-Protein-Coupled Lysophosphatidic Acid Receptors and Their Regulation of AKT Signaling. Int. J. Mol. Sci. 17, 215 (2016).

48. Joung, J. et al. Genome-scale activation screen identifies a lncRNA locus regulating a gene neighbourhood. Nature 548, 343–346 (2017).

49. Nogueira, T. C. et al. GLIS3, a susceptibility gene for type 1 and type 2 diabetes, modulates pancreatic beta cell apoptosis via regulation of a splice variant of the BH3-only protein Bim. PLoS Genet. 9, e1003532 (2013).

50. Rotem, A. et al. Alternative to the soft-agar assay that permits high-throughput drug and genetic screens for cellular transformation. Proc. Natl. Acad. Sci. U. S. A. 112, 5708–5713 (2015).

51. Yang, X. et al. A public genome-scale lentiviral expression library of human ORFs. Nat. Methods 8, 659–661 (2011).

52. Sack, L. M. et al. Profound Tissue Specificity in Proliferation Control Underlies Cancer Drivers and Aneuploidy Patterns. Cell 173, 499–514.e23 (2018).

53. Manguso, R. T. et al. In vivo CRISPR screening identifies Ptpn2 as a cancer immunotherapy target. Nature 547, 413–418 (2017).

54. Orchard, R. et al. Identification of anti-norovirus genes in mouse and human cells using genome-wide CRISPR activation screening. bioRxiv 350090 (2018). doi:10.1101/350090

55. Kiani, S. et al. Cas9 gRNA engineering for genome editing, activation and repression. Nat. Methods 12, 1051–1054 (2015).

56. Liao, H.-K. et al. In Vivo Target Gene Activation via CRISPR/Cas9-Mediated Trans-epigenetic Modulation. Cell 171, 1495–1507.e15 (2017).

55. Liao, H.-K. et al. In Vivo Target Gene Activation via CRISPR/Cas9-Mediated Trans-epigenetic Modulation. Cell 171, 1495–1507.e15 (2017).

